# Tryptophan and IFN-γ Differentially Modulate Cellular Uptake, Intracellular Trafficking, and Gene Expression of Messenger RNA-Loaded Lipid Nanoparticles in Dendritic Cells

**DOI:** 10.1101/2025.07.10.664160

**Authors:** Zitao Ma, Reena Jatyan, Nikki Crowley, Cierra Girardi, Mariya Khan, Akeile Smith, Pu-Sheng Wei, Yu-Hsuan Tsai, Kuo-Ching Mei

**Affiliations:** Department of Pharmaceutical Sciences, School of Pharmacy and Pharmaceutical Sciences, State University of New York at Binghamton, Johnson City, NY 13790, USA; Department of Molecular Pharmaceutics, College of Pharmacy, University of Utah, Salt Lake City, UT 84112, USA

**Author notes:** Equal Contribution (Zitao Ma, Reena Jatyan, Pu-Sheng Wei). Equal Contribution: Authors Listed Alphabetically by Last Names (Nikki Crowley, Cierra Girardi, Mariya Khan, Akeile Smith).

**Keywords:** messenger RNA lipid nanoparticles, dendritic cells, tryptophan deprivation, interferon-gamma stimulation, endosomal escape, immune tolerance

## Abstract

Lipid nanoparticle (LNP)-mediated mRNA delivery has emerged as a powerful platform for both immunostimulatory and immunomodulatory applications. However, the influence of local immunometabolic cues on the intracellular fate and translational efficiency of mRNA-LNPs remains poorly understood. In this study, we investigated how interferon-gamma (IFN-γ), a potent inducer of indoleamine 2,3-dioxygenase 1 (IDO1), and tryptophan (Trp) deprivation independently and combinatorially affect mRNA-LNP function in DC2.4 dendritic cells. These two cues are canonical drivers of immunoregulatory microenvironments, particularly those that favor tolerogenic dendritic cell programming and the induction of regulatory T cells. Using dual-reporter mRNA constructs and high-resolution confocal imaging, we show that IFN-γ stimulation reduces total cellular mRNA uptake and lysosomal accumulation without affecting translation efficiency and endosomal escape efficiency. Whereas Trp deprivation also reduces the overall cellular uptake of mRNA-LNPs, it also significantly impairs protein synthesis from mRNA-LNPs and modestly reduces endosomal escape, despite having minimal impact on lysosomal mRNA levels. Spatial compartmentalization analysis revealed that IFN-γ and Trp limitation disrupt distinct steps in the delivery-translation cascade, acting independently but additively to suppress the ultimate protein translation from mRNA-LNPs in DC2.4 dendritic cells. These findings highlight the importance of considering local metabolic and cytokine contexts when deploying mRNA-LNPs for immunological applications. Our work provides mechanistic insights into how immunoregulatory environments impair the delivery and translation of mRNA-LNPs, suggesting strategies to tune delivery outcomes for tolerogenic purposes.

## Introduction

The clinical approval and continued development of mRNA-lipid nanoparticle (mRNA-LNP) platforms have established a new paradigm in immunotherapy. Unlike traditional vaccines, mRNA-LNPs deliver messenger RNA into host cells, where it is translated into full-length antigenic proteins in the cytoplasm ^1^. The ability of mRNA-LNPs to elicit innate and adaptive immune responses is strongly influenced by their cellular uptake efficiency, endosomal escape, and intracellular processing in antigen-presenting cells, particularly dendritic cells. At the injection site, resident and recruited (monocyte-derived) dendritic cells internalize mRNA-LNPs, translate the mRNA into antigenic proteins, and process these antigens into peptide fragments (epitopes) that are presented on major histocompatibility complex (MHC) molecules to initiate T cell activation. This process supports both CD8⁺ cytotoxic T lymphocyte responses, via MHC class I presentation of endogenously synthesized antigens (e.g., cytosolic loading) ^2^, and CD4⁺ helper T cell responses, via MHC class II presentation of exogenous antigens (e.g., through phagocytic endosomal-lysosomal processing of secreted or cell-associated proteins) ^3^, or endogenous antigens delivered to lysosomes through autophagy ^4,5^.

While mRNA-LNP platforms have shown efficacy in eliciting immunogenic responses that can be further modulated by the choice of lipid formulations ^6,7^, they have also been explored as tolerogenic platforms to induce immune suppression in models of autoimmunity, allergy, and anaphylaxis by inducing regulatory T cell (Treg) responses ^8–11^. This could be strengthened by incorporating anti-inflammatory lipid drug conjugates, e.g., dexamethasomes ^7^, or rapamycin ^12^. However, little is known about the intracellular fate of mRNA-LNPs in dendritic cells under immune-suppressive or tolerogenic environments. To investigate this, we focused on one specific nature-derived pro-Treg environment, i.e., the tumor microenvironment (TME), where dendritic cells are frequently exposed to metabolic and cytokine-derived tolerogenic cues. Among these, tryptophan catabolism, driven by interferon-gamma (IFN-γ)-inducible enzyme indoleamine-2,3-dioxygenase 1 (IDO1), plays a critical immunoregulatory role ^13^. IDO1 depletes extracellular tryptophan while generating immunosuppressive kynurenine metabolites, thereby creating a microenvironment that inhibits effector T cells and favors the expansion and stability of Tregs ^14,15^. The IFN-γ-IDO1 axis thus constitutes a non-classical immune checkpoint, functionally analogous to PD-L1 and CTLA-4.

Most mechanistic studies on mRNA-LNP delivery have been conducted under standard culture conditions, lacking the metabolic and inflammatory complexity of tissue environments. In contrast, metabolic and inflammatory cues reprogram dendritic cell function, shaping the outcome of T-cell priming toward either immunogenic or tolerogenic responses depending on the context. These conditions may alter DC function by modulating protein synthesis, promoting autophagic flux, and disrupting canonical endosomal trafficking, all processes with direct implications for the intracellular fate of mRNA cargo. Despite growing interest in developing mRNA-LNPs for immunotherapy, the mechanisms by which IFN-γ and tryptophan deprivation influence the intracellular processing of mRNA-LNPs in dendritic cells remain poorly understood.

To address this gap, we investigated how tryptophan deprivation and IFN-γ stimulation modulate the intracellular processing of mRNA-LNPs in DC2.4 cells, a murine dendritic cell line with preserved antigen presentation function ^16^. Cells were cultured under four defined conditions representing increasing immunosuppressive pressure: tryptophan-replete, tryptophan-replete with IFN-γ, tryptophan-deprived, and tryptophan-deprived with IFN-γ. These experimental conditions model distinct immune environments, culminating in a cytokine- and metabolite-rich state that recapitulates features of a well-characterized tolerogenic context. We formulated mRNA-LNPs using SM-102^17^, a clinically validated ionizable lipid, and loaded them with luciferase mRNA for quantitative protein expression and Cy5-labeled eGFP mRNA for imaging-based tracking. We assessed cell viability and biosynthetic activity (MTT, CCK-8^18^, ATP, LDH^19^ assays), translation efficiency (luciferase assay), and intracellular trafficking (confocal microscopy). Together, these studies provide a mechanistic framework to evaluate how immune context reshapes dendritic cell handling of mRNA-LNPs, offering foundational insights into how immune context governs the intracellular fate of mRNA-LNPs in antigen-presenting cells.

## Materials and Methods

All reagents and materials used in this study are detailed in the Supporting Information. Briefly, the mouse dendritic cell line DC2.4 was maintained in RPMI-1640-based media under standard cell culture conditions. Experimental treatments involved modulating tryptophan concentrations (100% or 50%) and IFN-γ supplementation (100 ng/mL) to mimic distinct immunoregulatory environments. mRNA-loaded lipid nanoparticles (mRNA-LNPs) were formulated using an ethanol injection method with a defined lipid composition (SM-102/DSPC/cholesterol/DMG-PEG2kDa, molar ratio 50:10:38.5:1.5) and characterized by dynamic light scattering and RiboGreen assay. Cy5-labeled eGFP mRNA and firefly luciferase (FLuc) mRNA were used for intracellular tracking and protein expression assays, respectively. Cellular viability and metabolic responses were assessed using MTT, CCK-8, LDH, and ATP luminescence assays. Gene expression and endosomal escape were evaluated by FLuc quantification and live-cell confocal microscopy following treatment with eGFP-Cy5 mRNA-LNPs and LysoTracker staining. Fluorescence intensity measurements were quantified using ImageJ, and all statistical analyses were performed using GraphPad Prism 10. Comprehensive experimental procedures, media compositions, reagent sources, and data acquisition methods are available in the online Supporting Information.

## Results

### Baseline characterization of dendritic cell viability and metabolism under tryptophan deprivation and IFN-γ stimulation

To evaluate how nutritional stress and inflammatory signaling jointly affect the viability and metabolism of dendritic cells, we systematically profiled DC2.4 cells exposed to combinatorial conditions of tryptophan (Trp) deprivation and interferon-gamma (IFN-γ) stimulation. As illustrated in the experimental workflow (**Fig. 1a**), DC2.4 cells were first allowed to adhere and stabilize for 12 hours in standard DC2.4 cell culture medium, followed by a 36-hour treatment period under one of four conditions: (1) 100% Trp (control), (2) 100% Trp + IFN-γ (100 ng/mL), (3) 50% Trp (Trp deprivation or Trp-Dep in short), or (4) 50% Trp + IFN-γ. These conditions model a range of physiological and pathological stress scenarios, including nutrient-limited microenvironments and immune activation. We employed four complementary assays to capture distinct aspects of cell health and metabolism (**Fig. 1b**). The LDH assay quantifies extracellular lactate dehydrogenase as a marker of plasma membrane rupture, while the MTT assay reflects mitochondrial reductive capacity. The ATP assay measures intracellular energy content via luminescence, and the CCK-8 assay detects cytosolic NAD(P)H-dependent dehydrogenase activity, offering insights into the redox status of the cytoplasm.

**Figure 1.**
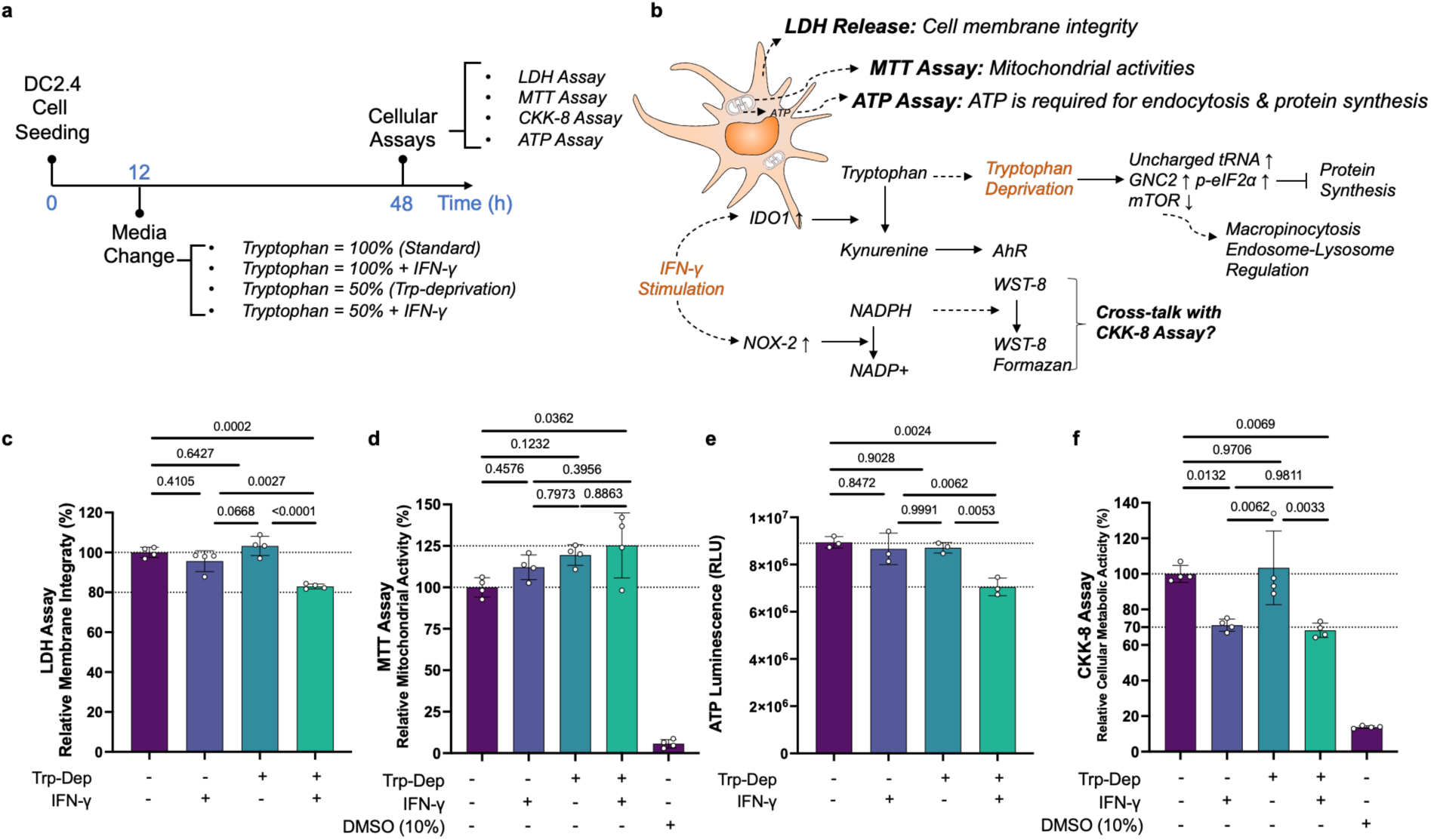
Baseline characterization of DC2.4 cell viability, mitochondrial activity, and ATP levels under combinatorial conditions of IFN-γ stimulation and tryptophan deprivation. **(a)** Schematic experimental workflow: DC2.4 dendritic cells were seeded and pre-incubated for 12 h, followed by 36 h of exposure to one of four treatment conditions combining 100% or 50% tryptophan (Trp vs. Trp-Deprivation) with or without 100 ng/mL IFN-γ stimulation, prior to viability and metabolic assays. **(b)** Overview of assay targets: LDH assay reflects cell membrane integrity; MTT assay reports on mitochondrial reductive activity; ATP assay quantifies intracellular ATP; and CCK-8 reports cytosolic NAD(P)H-dependent dehydrogenase activity. **(c)** LDH assay revealed significantly elevated membrane damage only in the dual-treatment group (50% Trp-Dep + IFN-γ), with a significant interaction: IFN-γ, F(1,12) = 41.14, *p* < 0.0001; Trp deprivation, F(1,12) = 6.02, *p* = 0.0304; interaction, F(1,12) = 17.14, *p* = 0.0014. **(d)** MTT assay showed increased mitochondrial activity under all stress conditions, especially with Trp deprivation alone (19.5% increase vs. control, F(1,12) = 8.28, *p* = 0.0139), with no significant interaction (F(1,12) = 0.32, *p* = 0.5842). DMSO control confirmed assay validity. **(e)** ATP levels decreased significantly only under combined Trp deprivation and IFN-γ stimulation, with significant main effects and interaction: IFN-γ, F(1,8) = 16.27, *p* = 0.0038; Trp, F(1,8) = 14.77, *p* = 0.0049; interaction, F(1,8) = 8.32, *p* = 0.0204. **(f)** CCK-8 assay showed a pronounced decrease in cytosolic metabolic activity upon IFN-γ stimulation under both Trp conditions (−28.9% at 100% Trp; −34.3% at 50% Trp), independent of Trp availability (IFN-γ, F(1,12) = 34.22, *p* < 0.0001; no Trp or interaction effect). Despite a reduced signal, LDH, MTT, and ATP assays did not corroborate cytotoxicity, suggesting that the CCK-8 reduction arises from IFN-γ-induced NOX2 activation, which depletes cytosolic NADPH, the electron donor necessary for tetrazolium dye reduction. DMSO toxicity controls confirmed CCK-8 assay performance. All together, these data demonstrate that IFN-γ and Trp deprivation synergistically compromise membrane integrity and ATP homeostasis. The mitochondrial reductive function was elevated under Trp deprivation and IFN-γ stimulation, while cytosolic dehydrogenase activity (CCK-8) is selectively suppressed under IFN-γ stimulation. This suggests that localized redox depletion, rather than global metabolic failure, is a confounding factor in interpreting CCK-8 readouts under inflammatory conditions.

Cell membrane integrity was first assessed by quantifying LDH release (**Fig. 1c**). Minimal LDH signal was observed in cells under single-stressor conditions (either Trp deprivation or IFN-γ alone), indicating preserved membrane integrity. However, when cells were simultaneously exposed to both IFN-γ and reduced Trp, LDH release increased significantly, suggesting compromised membrane integrity under dual stress. Two-way ANOVA confirmed a highly significant main effect of IFN-γ (*F*(1,12) = 41.14, *p* < 0.0001) and a smaller but significant effect of Trp deprivation (*F*(1,12) = 6.02, *p* = 0.0304). Importantly, a significant interaction between the two factors (*F*(1,12) = 17.14, *p* = 0.0014) was detected, supporting a synergistic effect in inducing membrane damage.

Mitochondrial reductive activity, as evaluated by the MTT assay (**Fig. 1d**), displayed a distinct trend. Surprisingly, all three stress conditions—including IFN-γ stimulation, Trp deprivation, and their combination—elicited elevated MTT signals relative to the untreated control. The effect was most pronounced in the Trp deprivation alone group, which exhibited a 19.5% increase in MTT activity compared to control. Statistical analysis confirmed a significant main effect of Trp deprivation (*F*(1,12) = 8.28, *p* = 0.0139), while the effect of IFN-γ was not statistically significant (*F*(1,12) = 2.66, *p* = 0.1293), and no interaction was observed (*F*(1,12) = 0.32, *p* = 0.5842). These results suggest that mitochondrial function, rather than being suppressed, may be upregulated under nutrient limitation, possibly as a compensatory adaptation.

Intracellular ATP content, measured by luminescence assay, provided a sensitive readout of cellular energy status (**Fig. 1e**). Cells maintained ATP levels under single-stressor conditions but exhibited a significant ATP depletion only when both IFN-γ stimulation and Trp deprivation were applied simultaneously. This pattern suggests a threshold effect, where dual stress disrupts bioenergetic homeostasis. Two-way ANOVA revealed significant main effects of both IFN-γ (*F*(1,8) = 16.27, *p* = 0.0038) and Trp deprivation (*F*(1,8) = 14.77, *p* = 0.0049), along with a significant interaction (*F*(1,8) = 8.32, *p* = 0.0204), further supporting a synergistic energy stress phenotype.

Cytosolic redox activity, as determined by the CCK-8 assay (**Fig. 1f**), was markedly and consistently suppressed by IFN-γ stimulation, regardless of Trp concentration. Specifically, IFN-γ reduced CCK-8 signal by 28.9% under 100% Trp and 34.3% under 50% Trp conditions. Two-way ANOVA confirmed a highly significant main effect of IFN-γ (*F*(1,12) = 34.22, *p* < 0.0001), while neither Trp deprivation (*F*(1,12) = 0.3561, *p* = 0.5614) nor the interaction term (*F*(1,12) = 0.07723, *p* = 0.7859) was significant. Importantly, this suppression occurred in the absence of elevated LDH release or decreased ATP, suggesting that it does not reflect overt cytotoxicity. Rather, the CKK-8 assay was likely cross-talking with IFN-γ-induced upregulation of NADPH oxidase (NOX2), which consumes cytosolic NADPH required for CCK-8 dye reduction, thereby artificially lowering the readout signal.

Together, these data establish a nuanced baseline understanding of DC2.4 cell responses under metabolic and inflammatory stress. While Trp deprivation and IFN-γ stimulation exert modest effects when applied independently, their combination leads to synergistic disruption of membrane integrity and ATP homeostasis. Mitochondrial reductive activity remains preserved or elevated, whereas CCK-8 signal is selectively impaired by IFN-γ, likely due to localized NADPH depletion rather than global metabolic failure. These findings underscore the importance of integrating multiple metabolic readouts to accurately interpret cellular stress responses in dendritic cells.

### Physicochemical characterization of mRNA-loaded lipid nanoparticles (LNPs)

To enable downstream investigation of mRNA uptake, intracellular trafficking, and translation under defined immunometabolic conditions, we formulated three LNP formulations, each encapsulating a specific mRNA cargo. These included: (1) Firefly luciferase mRNA (Fluc-mRNA-LNPs), to be used for luminescence-based quantification of protein translation; (2) Cy5-labeled mRNA encoding enhanced green fluorescent protein (eGFP-Cy5-mRNA-LNPs), designed to enable dual-mode imaging of mRNA uptake (via Cy5) and protein expression (via eGFP) in live-cell confocal microscopy studies; (3) eGFP-scrambled-mRNA-LNP, a translation-incompetent RNA construct without Cy5 labeling, serving as a dual-negative control for confocal microscopic imaging analyses.

All three LNPs were prepared using a rapid ethanol injection method published previously ^7,20^. The formulation was based on a clinically validated ionizable lipid formulation composed of SM-102, DSPC, cholesterol, and DMG-PEG2000 at a molar ratio of 50:10:38.5:1.5 ^1^. Assembly was performed by mixing lipids dissolved in ethanol with aqueous mRNA in citric acid buffer (pH 4.0), at an NP ratio of 1:6 and a volume ratio of 1:3 (v/v). This approach ensures electrostatic complexation of mRNA with ionizable lipids under acidic conditions, followed by nanoparticle formation upon dilution. Formulations were immediately diluted, filtered, and stored at 4°C for short-term use (<1 day) to minimize degradation and aggregation. To ensure equivalent mRNA dosing across conditions, RiboGreen-based encapsulation assays were performed to determine mRNA loading. This allowed all cell-based experiments to be conducted at a normalized dose of 200 ng mRNA per well, avoiding confounding effects due to variation in RNA loading or delivery input.

We next performed standard biophysical quality control on all mRNA-LNP formulations, including dynamic light scattering (DLS), polydispersity index (PDI), and zeta potential measurements (**Fig. 2a–c**). DLS analysis revealed mean hydrodynamic diameters ranging from 125.4 to 153.0 nm across the four formulations (**Fig. 2a**). One-way ANOVA detected significant variation among formulations (F(3,8) = 6.749, *p* = 0.0139). Notably, post hoc Tukey’s test showed that the eGFP-Cy5-mRNA-LNPs were significantly smaller in diameter than both Fluc-mRNA-LNPs (*p* = 0.0190) and eGFP-scrambled-mRNA-LNPs (p = 0.0196), with no other pairwise differences reaching significance. This size reduction may be attributed to Cy5 conjugation altering mRNA conformation or electrostatic interactions during self-assembly, potentially leading to more compact particles.

**Figure 2.**
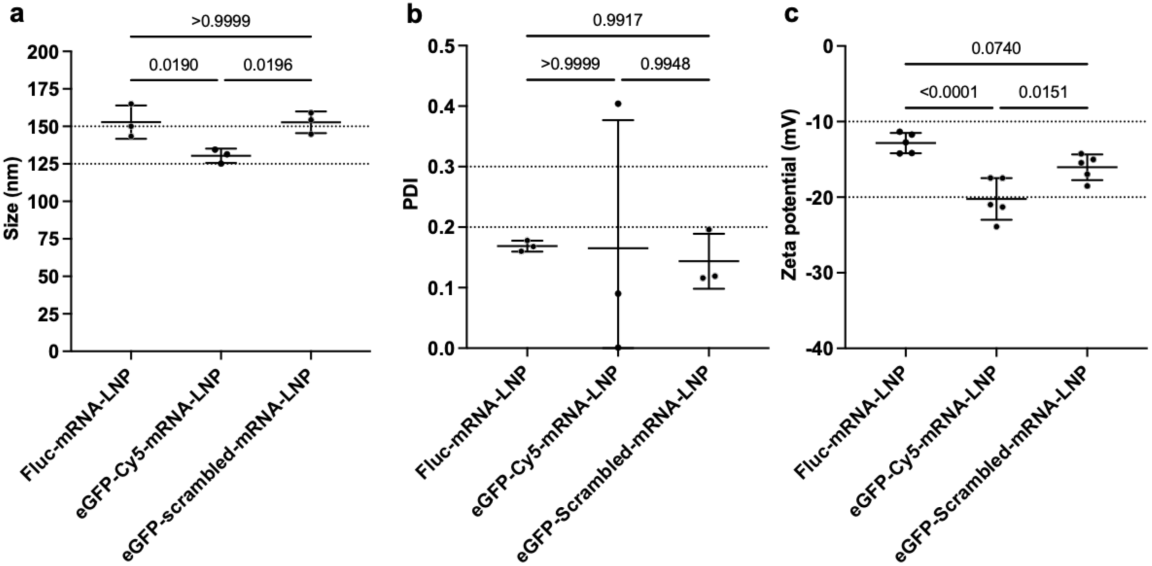
Physicochemical characterization of four mRNA-LNP formulations used for downstream expression and imaging studies. This figure summarizes the quality control and biophysical characterization of four lipid nanoparticle (LNP) formulations encapsulating different mRNA cargoes: (1) Firefly luciferase mRNA (Fluc-mRNA-LNP), used for luminescence-based transfection quantification; (2) Cy5-labeled eGFP mRNA (eGFP-Cy5-mRNA), designed for dual tracking of uptake (Cy5) and translation (eGFP) in confocal microscopy; (3) eGFP-scrambled-mRNA, containing translation-incompetent, unlabeled RNA to serve as a negative control for both Cy5 and eGFP imaging. **(a)** Dynamic light scattering (DLS) revealed mean hydrodynamic diameters ranging from ∼125 to 153 nm. One-way ANOVA indicated significant differences across formulations (F(3,8) = 6.749, *p* = 0.0139). Tukey’s multiple comparisons test showed that eGFP-Cy5-mRNA-LNPs were significantly smaller than Fluc-mRNA-LNPs (*p* = 0.0190) and eGFP-scrambled-mRNA-LNPs (*p* = 0.0196). No other size differences were statistically significant. **(b)** Polydispersity index (PDI) values were generally <0.3, indicating acceptable uniformity. One outlier batch of eGFP-Cy5-mRNA-LNP showed a higher PDI, but overall group differences were not significant (F(3,8) = 0.031, *p* = 0.9921), confirming consistent formulation quality across groups. **(c)** Zeta potential measurements showed all formulations carried anionic surface charges ranging from −12.8 to −20.2 mV. One-way ANOVA showed significant variation across groups (F(3,16) = 14.82, *p* < 0.0001). eGFP-Cy5-mRNA-LNP had a significantly more negative zeta potential than Fluc-mRNA-LNP (*p* < 0.0001) and eGFP-scrambled-mRNA-LNP (*p* = 0.0151), likely due to Cy5 conjugation altering surface charge properties. Collectively, these data confirm that all mRNA-LNPs fall within acceptable size and charge ranges for in vitro application. The physicochemical distinctions observed with Cy5-labeled RNA, specifically its smaller size and increased surface negativity, should be carefully considered when interpreting trafficking and expression results in downstream cellular studies.

PDI analysis confirmed that all LNPs exhibited acceptable colloidal uniformity, with values consistently below the 0.3 threshold commonly used to define monodisperse preparations (**Fig. 2b**). One formulation batch of eGFP-Cy5-mRNA-LNPs (out of three independent batches) showed a slightly elevated PDI, which did not exceed 0.35 and remained functionally acceptable. Statistically, there were no significant differences in PDI across groups (F(3,8) = 0.031, p = 0.9921), indicating high reproducibility and consistency in formulation.

Zeta potential analysis revealed that all LNPs carried net negative surface charges, with values ranging from −12.8 to −20.2 mV (**Fig. 2c**), consistent with partial PEGylation and surface-exposed phosphate residues of the mRNA. However, significant variation in surface charge was detected (F(3,16) = 14.82, *p* < 0.0001), driven primarily by the eGFP-Cy5-mRNA-LNPs. Compared to the Fluc-mRNA-LNPs and scrambled-mRNA-LNPs, the Cy5-labeled formulation displayed significantly more negative zeta potential (*p* < 0.0001 and *p* = 0.0151, respectively). This suggests that fluorescent labeling may introduce additional anionic moieties or alter the spatial configuration of surface-associated mRNA, impacting the surface electrostatic profile. Importantly, these biophysical deviations might need to be considered when interpreting differences in intracellular trafficking or endosomal escape across formulations, particularly in imaging studies that rely on Cy5-labeled RNA.

In sum, all mRNA-LNP formulations were successfully prepared with acceptable size distribution and physiologically relevant surface charge. While Cy5 labeling introduced modest yet statistically significant differences in hydrodynamic size and zeta potential, the overall properties remained within the established range for efficient cellular delivery. These preparations provide a well-controlled toolkit for probing mRNA delivery and translation under diverse immunometabolic perturbations in dendritic cells.

### Tryptophan Deprivation and IFN-γ Synergistically Impair mRNA-LNP-Mediated Expression in Dendritic Cells

To investigate how inflammatory and metabolic stresses resulting from tryptophan deprivation impact the functional delivery and expression of exogenous mRNA via lipid nanoparticles (LNPs), we employed a combinatorial stress model using DC2.4 dendritic cells as mentioned previously. Cells were pretreated for 12 hours with either full-nutrient media (100% tryptophan), 50% tryptophan-deprived media, or the same media supplemented with interferon-gamma (IFN-γ, 100 ng/mL), followed by treatment with mRNA-loaded LNPs (**Fig. 3a**). Firefly luciferase mRNA (Fluc-mRNA-LNP) was used to quantify translation efficiency and evaluate possible cytotoxicity resulting from the LNPs, while Cy5-labeled eGFP mRNA (eGFP-Cy5-mRNA-LNP) was employed for imaging of uptake and intracellular protein expression.

**Figure 3.**
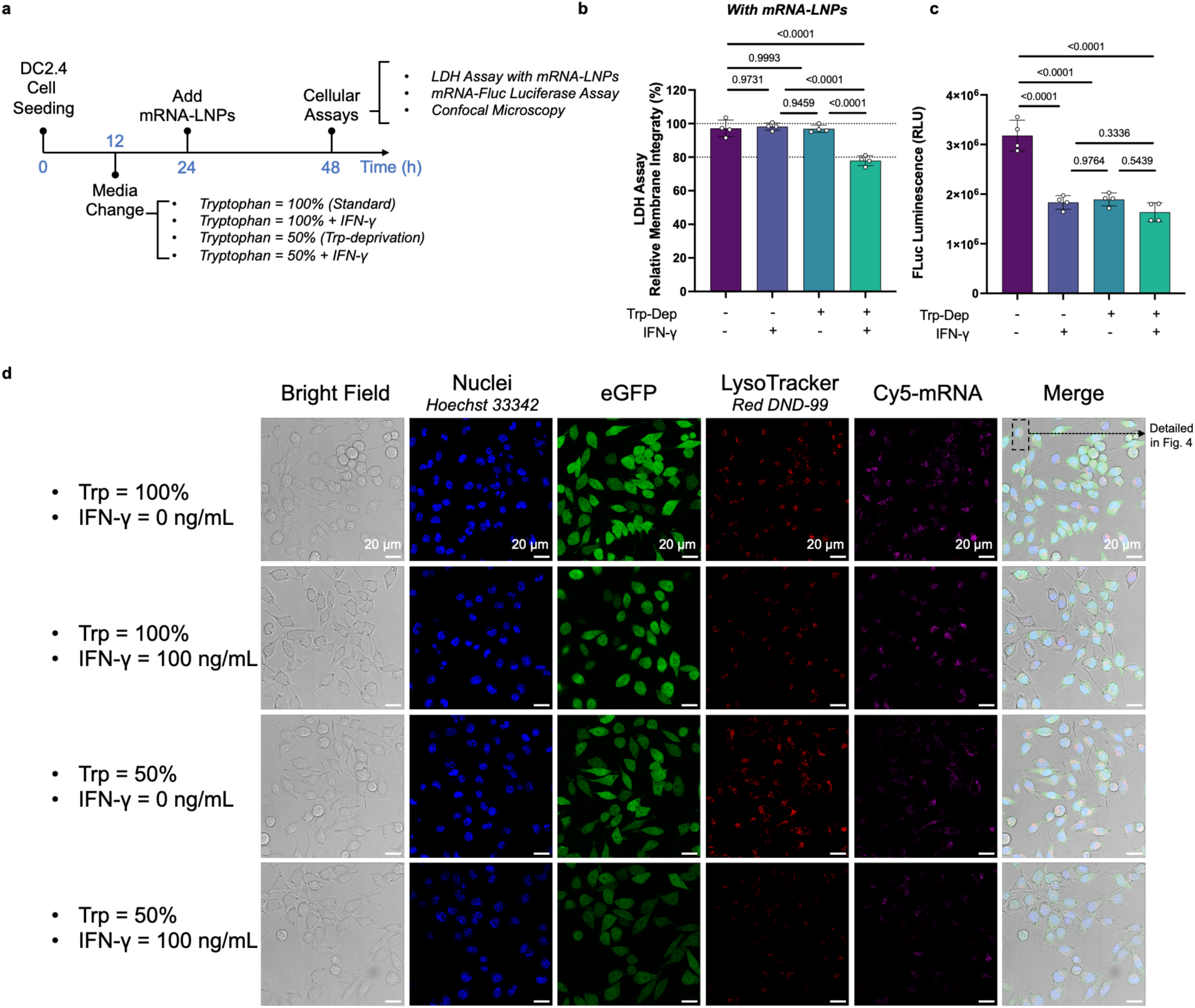
Effects of Tryptophan Deprivation and IFN-γ Stimulation on mRNA-LNP-Induced Membrane Integrity, Translation Efficiency, and Intracellular Trafficking in DC2.4 Cells. **(a)** Experimental timeline. DC2.4 cells were seeded and cultured for 12 h before switching to four treatment conditions: 100% tryptophan (Trp) or 50% Trp deprivation, with or without IFN-γ (100 ng/mL). At 24 h post-seeding, cells were treated with Fluc-mRNA-LNPs (for LDH and luciferase assays) or eGFP-Cy5-mRNA-LNPs / eGFP-scrambled-mRNA-LNPs (for imaging). Assays were conducted at 48 h. **(b)** LDH assay indicates that mRNA-LNP treatment does not independently disrupt membrane integrity. However, dual treatment (50% Trp + IFN-γ) resulted in significant LDH release. Two-way ANOVA confirmed main effects of IFN-γ (F(1,12) = 30.81, *p*= 0.0001), Trp deprivation (F(1,12) = 39.97, *p* < 0.0001), and their interaction (F(1,12) = 37.84, *p* < 0.0001), consistent with prior observations (Fig. 1c). **(c)** Firefly luciferase assay, normalized to LDH to control for cell loss, revealed that both IFN-γ and Trp deprivation suppress Fluc-mRNA-LNP-mediated Fluc protein expression. Two-way ANOVA showed significant main effects of IFN-γ (F(1,12) = 60.72, *p* < 0.0001), Trp deprivation (F(1,12) = 52.11, *p* < 0.0001), and their interaction (F(1,12) = 28.00, *p* = 0.0002). IFN-γ reduced luciferase expression by 42.1% (100% Trp) and 30.6% (50% Trp), while Trp deprivation alone reduced expression by 40.8% (no IFN-γ) and 29.1% (with IFN-γ). **(d)** Confocal imaging of eGFP-Cy5-mRNA-LNP-treated DC2.4 cells was used to visualize mRNA uptake (Cy5), lysosomal localization (LysoTracker), and protein expression (eGFP). Cells were co-stained with Hoechst 33342 for nuclei. The negative control (scrambled-mRNA-LNP, not shown) confirmed the background signal. Under full nutrient and unstimulated conditions, robust Cy5 and eGFP signals were observed. Upon IFN-γ stimulation, Trp deprivation, or their combination, eGFP expression was markedly reduced. Cy5 signal also appeared diminished under all stress conditions, suggesting that uptake may be partially compromised. These findings demonstrate that IFN-γ and Trp deprivation impair mRNA-LNP-mediated protein expression at multiple levels, including potential effects on uptake and intracellular processing, with translational suppression being a consistent outcome under stress.

As previously introduced, cell membrane integrity, as measured by lactate dehydrogenase (LDH) release, was largely unaffected by LNP treatment alone. The overall result is mirroring the prior study without mRNA-LNP treatment shown in **Fig. 1c** and **Fig. 3b**, i.e., under dual-stress conditions (50% Trp deprivation combined with IFN-γ stimulation), a significant increase in LDH release was observed (**Fig. 3b**). Two-way ANOVA revealed significant main effects of IFN-γ (F(1,12) = 30.81, p = 0.0001), tryptophan deprivation (F(1,12) = 39.97, *p* < 0.0001), and a strong interaction between the two (F(1,12) = 37.84, *p* < 0.0001), indicating that the combination of metabolic and inflammatory stimuli compromises cell viability or membrane integrity.

Firefly luciferase expression, normalized to LDH levels to account for cell loss, was significantly reduced across all stressed conditions (**Fig. 3c**). Two-way ANOVA confirmed significant main effects of IFN-γ (F(1,12) = 60.72, *p* < 0.0001), Trp deprivation (F(1,12) = 52.11, *p* < 0.0001), and their interaction (F(1,12) = 28.00, *p* = 0.0002). IFN-γ treatment alone reduced luciferase output by 42.1% in full-nutrient media and by 30.6% in Trp-deprived conditions. Trp deprivation in the absence of IFN-γ led to a 40.8% decrease in expression, while its presence resulted in a 29.1% reduction. These data indicate that both immunological and metabolic stressors independently and synergistically suppress the translation of exogenous mRNA.

To further dissect the mechanistic basis of reduced protein output, we performed live-cell confocal imaging following treatment with eGFP-Cy5-mRNA-LNPs. Under non-stressed conditions, robust Cy5 fluorescence (tracking cellular uptake and intracellular localization of the mRNA) and eGFP expression (reflecting protein translation from the mRNA) were observed in the cytoplasm of DC2.4 cells (**Fig. 3d**). In contrast, both IFN-γ stimulation and Trp deprivation led to marked reductions in eGFP signal, with the most pronounced suppression observed under combined stress. A concurrent decrease in Cy5 fluorescence was also noted, suggesting impaired uptake or altered intracellular trafficking of the LNPs. The absence of signal in cells treated with translation-incompetent eGFP-scrambled-mRNA-LNPs (negative control, not shown) confirmed the specificity of both fluorescence channels.

Collectively, these data demonstrate that IFN-γ and tryptophan deprivation compromise mRNA-LNP-mediated protein expression in DC2.4 cells through multiple mechanisms, including potential reductions in cellular uptake and clear suppression of translational efficiency. These findings underscore the importance of immunometabolic context when interpreting the efficacy of non-viral mRNA delivery systems in antigen-presenting cells.

### Single-Cell Imaging Reveals Additive Impairments in mRNA Uptake and Translation by IFN-γ and Tryptophan Deprivation

Representative confocal images of cells under control conditions (100% Trp, no IFN-γ) show distinct cytoplasmic fluorescence in both channels (**Fig. 4a**). Regions of interest (ROIs) were drawn around the cytoplasm of individual cells (*n* = 48 per group), and signal intensities were extracted and quantified across conditions. Nuclear staining with Hoechst was used to exclude overlapping or apoptotic cells, while Lysotracker (not shown) marked endo-lysosomal compartments for future trafficking analysis in **Fig. 5**.

**Figure 4.**
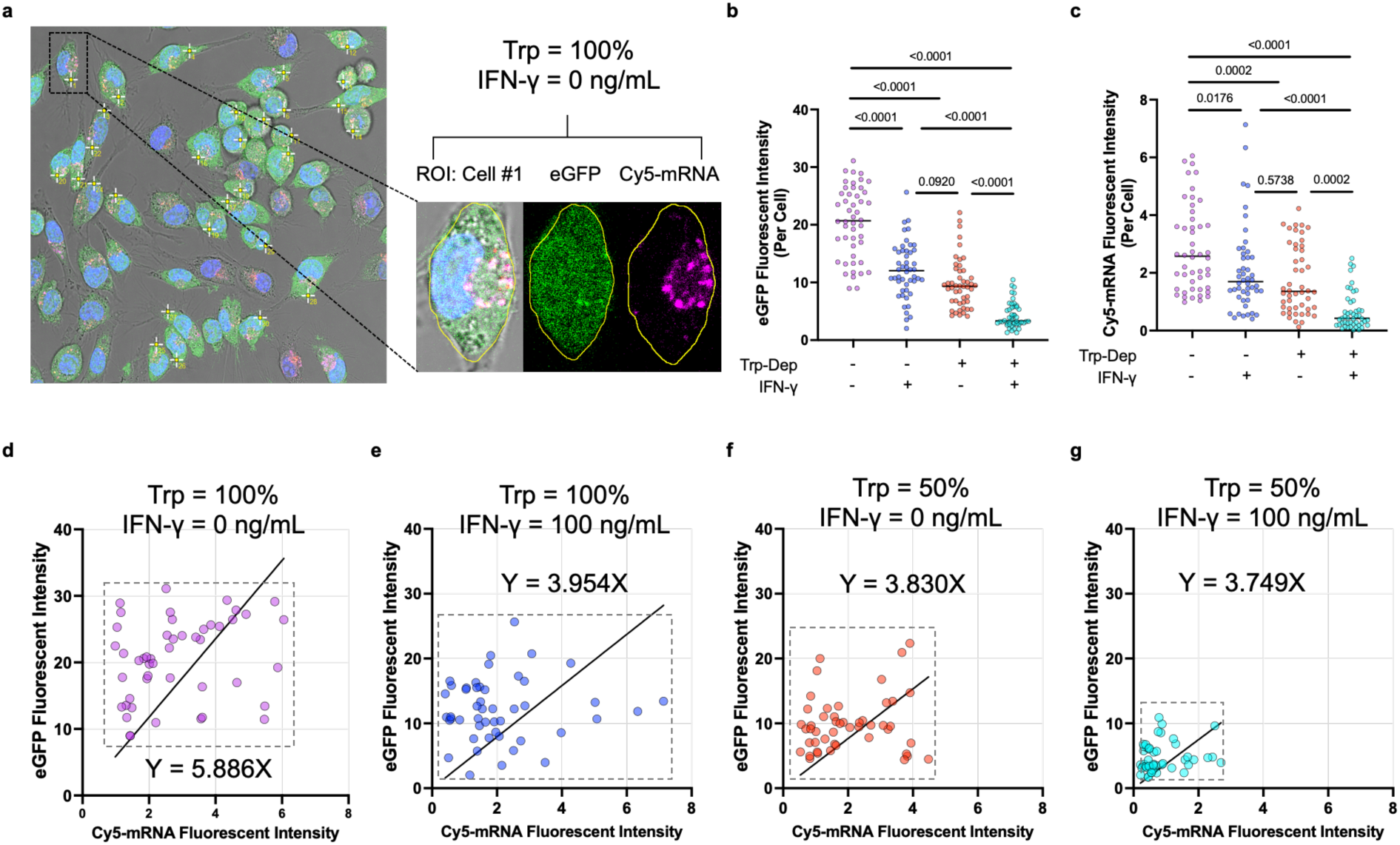
Quantitative single-cell imaging analysis of mRNA-LNP uptake and translation efficiency in DC2.4 cells under varying tryptophan availability and IFN-γ stimulation. **(a)** Representative 100× confocal image of DC2.4 cells cultured under 100% tryptophan without IFN-γ, corresponding to the control condition in Figure 3d. Individual cells were segmented into regions of interest (ROIs) based on cytoplasmic boundaries, enabling quantification of Cy5-eGFP-mRNA fluorescence (magenta; mRNA uptake) and eGFP expression (green; eGFP-mRNA translation). Two cells are enlarged to show signal extraction. Hoechst (blue) and Lysotracker Red (not shown here) were used for nuclear and lysosomal counterstaining. **(b)** Single-cell quantification of total eGFP signal (*n* = 48 cells per group) revealed significant reductions with both IFN-γ stimulation and Trp deprivation. Two-way ANOVA showed main effects of IFN-γ (F(1,188) = 104.6, *p* < 0.0001); Trp level (F(1,188) = 183.4, *p* < 0.0001). Trp level (F(1,188) = 104.6, *p* < 0.0001), and a significant interaction (F(1,188) = 5.454, *p* = 0.0206). Under 100% Trp, IFN-γ reduced eGFP intensity from 20.48 ± 6.14 to 12.14 ± 4.84. Under 50% Trp, IFN-γ further reduced eGFP intensity from 9.94 ± 4.31 to 4.69 ± 2.23. Trp deprivation reduced eGFP both with (12.14 → 4.69) and without (20.48 → 9.94) IFN-γ. These results demonstrate that IFN-γ and Trp deprivation independently impair translation, with additive suppressive effects. **(c)** Quantification of Cy5 signal per cell (*n* = 48 cells per group) indicated reduced mRNA uptake under both stressors. Two-way ANOVA revealed significant effects of IFN-γ (F(1,188) = 26.51, *p* < 0.0001) and Trp deprivation (F(1,188) = 48.52, *p* < 0.0001), without interaction (F(1,188) = 0.9001, *p* = 0.3440). IFN-γ reduced Cy5 from 2.84 ± 1.49 to 2.09 ± 1.49 under 100% Trp, and from 1.76 ± 1.18 to 0.66 ± 0.65 under 50% Trp. Trp deprivation reduced Cy5 both with (2.09 → 0.66) and without (2.84 → 1.76) IFN-γ. These findings confirm that both inflammatory and nutritional stressors reduce mRNA uptake, although their effects are not synergistic. **(d–g)** Correlation plots between Cy5 and eGFP intensities under the four conditions: **(d)** 100% Trp, no IFN-γ; **(e)** 100% Trp, +IFN-γ; **(f)** 50% Trp, no IFN-γ; **(g)** 50% Trp, +IFN-γ. Linear regression, constrained to the origin (Y = aX), was used to estimate translation efficiency per unit of mRNA uptake. The slope progressively decreased across (d) to (g), confirming reduced translation efficiency under IFN-γ, Trp deprivation, and their combination.

**Figure 5.**
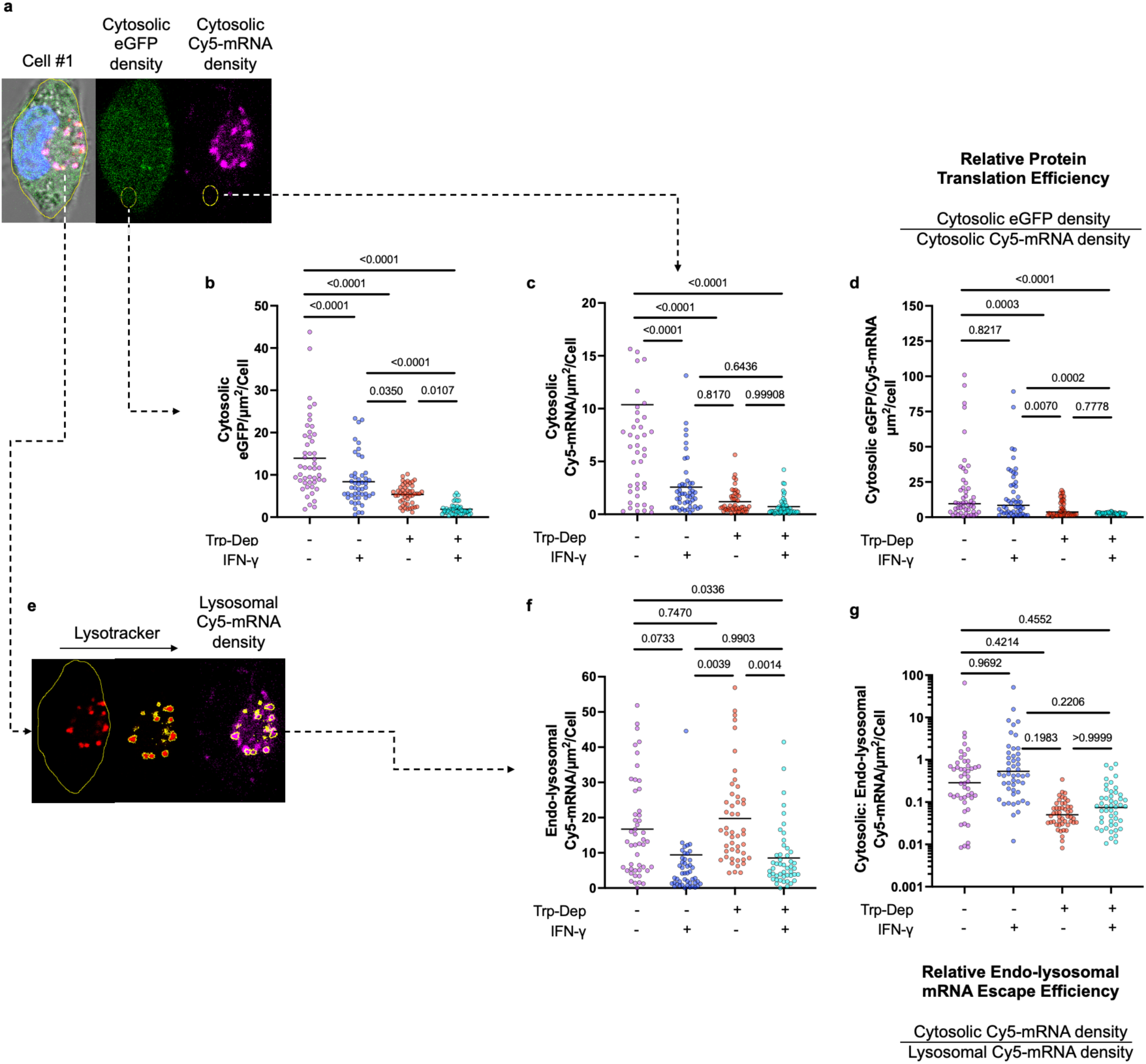
Spatial compartmentalization analysis reveals distinct effects of IFN-γ stimulation and tryptophan deprivation on mRNA localization, translation, and endosomal escape in DC2.4 cells. **(a)** Representative high-resolution confocal image (100×) of DC2.4 Cell #1 (same as Figure 4a) under 100% tryptophan without IFN-γ treatment. Six cytoplasmic ROIs (circled in yellow, one is shown) were selected per cell, excluding the nucleus and LysoTracker-positive compartments, and confined within the cell border. For each ROI, the ratio of eGFP signal (translated protein from mRNA) to Cy5 signal (endo-lysosomal escaped eGFP-mRNA) was calculated per surface area (µm²), and averaged across ROIs to yield a per-cell translation efficiency score. **(b)** Quantification of cytosolic eGFP intensity per µm² across four experimental groups (*n* = 48 cells per group). Two-way ANOVA revealed significant effects of tryptophan deprivation (*F*(1,188) = 92.82, *p* < 0.0001) and IFN-γ stimulation (*F*(1,188) = 33.39, *p* < 0.0001), with no interaction. Both factors independently suppressed cytoplasmic protein synthesis, with the most profound reduction under combined treatment. **(c)** Quantification of cytosolic mRNA-Cy5 signal per µm². Both tryptophan deprivation and IFN-γ significantly reduced cytosolic mRNA (*p* < 0.0001 and *p* = 0.0003, respectively), with a significant interaction (*p* = 0.0012), indicating that cytosolic mRNA availability is affected by both metabolic and inflammatory inputs. **(d)** Cytosolic translation efficiency (eGFP/mRNA-Cy5 per µm²). Two-way ANOVA revealed a significant main effect of tryptophan deprivation (*F*(1,188) = 34.91, *p* < 0.0001), but not of IFN-γ (*F*(1,188) = 1.651, *p* = 0.2004), nor an interaction (*F*(1,188) = 0.00342, *p* = 0.9534). Specifically, translation efficiency was reduced by 69.5% under 50% Trp (mean = 5.89) compared to 100% Trp (mean = 19.33) without IFN-γ, and by 83.0% under 50% Trp (mean = 2.8) vs. 100% Trp (mean = 16.51) with IFN-γ. **(e)** Lysosomal ROIs were defined by LysoTracker-positive compartments (magenta), and six per cell were selected and averaged to quantify lysosome-localized mRNA (Cy5) per µm². This spatial definition ensured the exclusion of cytosolic signals and allowed for the direct measurement of mRNA retention in endosomes/lysosomes. **(f)** Quantification of lysosomal mRNA signal (Cy5/µm²). Two-way ANOVA revealed a significant main effect of IFN-γ (*F*(1,188) = 19.13, *p* < 0.0001), but no significant effect of tryptophan deprivation (*F*(1,188) = 0.2461, *p* = 0.6204), nor interaction (*F*(1,188) = 0.8544, *p* = 0.3565). IFN-γ significantly reduced lysosomal mRNA density, suggesting decreased uptake or enhanced degradation. Post hoc analysis showed that the strongest drop occurred between 50% Trp ± IFN-γ (mean difference = 11.22, *p* = 0.0014), while tryptophan status alone had no significant impact. **(g)** Ratio of cytosolic to lysosomal mRNA signal (Cy5), used as a proxy for *endosomal escape efficiency.* Two-way ANOVA identified a significant main effect of Trp deprivation (*F*(1,188) = 5.974, *p* = 0.0154), with no significant effect of IFN-γ (*p* = 0.7197) or interaction (*p* = 0.7800). Despite the overall significance, multiple comparisons failed to detect a significant pairwise difference between individual groups. The data suggest a modest impairment of endosomal escape efficiency under tryptophan deprivation, but not under IFN-γ. In conclusion, Trp deprivation impairs both cytosolic mRNA localization and translation efficiency, consistent with global repression of protein synthesis. IFN-γ primarily reduces lysosomal mRNA content without affecting escape or translation efficiency. Together, these findings demonstrate that nutrient stress and inflammatory signaling impact distinct steps of mRNA processing and translation in dendritic cells.

We first looked at the total eGFP signal per cell, a proxy for successful mRNA translation, was significantly decreased, either under Trp-deprivation, or with IFN-γ stimulation, or both (**Fig. 4b**). Statistical analysis using two-way ANOVA revealed highly significant main effects of IFN-γ (F(1,188) = 104.6, *p* < 0.0001) and Trp deprivation (F(1,188) = 183.4, *p* < 0.0001), as well as a smaller but significant interaction (F(1,188) = 5.454, *p* = 0.0206), suggesting partially additive effects. Under full Trp conditions, IFN-γ reduced eGFP intensity from 20.48 ± 6.14 to 12.14 ± 4.84 per cell. Trp deprivation alone reduced eGFP expression by more than 50% (20.48 → 9.94), and the combination of both IFN-γ and 50% Trp led to the lowest observed expression level (4.69 ± 2.23), amounting to a 77.1% drop from baseline. These results suggest that both IFN-γ and Trp depletion independently impair the translational capacity of DC2.4 cells, likely via distinct mechanisms that converge on global mRNA utilization.

We next examined total Cy5 fluorescence signal per cell, which reflects the amount of mRNA delivered by LNPs and retained in each cell, independent of its translation (**Fig. 4c**). Both stressors again resulted in significant suppression. Two-way ANOVA revealed significant effects of IFN-γ (F(1,188) = 26.51, *p* < 0.0001) and Trp deprivation (F(1,188) = 48.52, *p* < 0.0001), but no significant interaction (F(1,188) = 0.9001, *p* = 0.3440), indicating that the impairments in mRNA uptake are additive but mechanistically independent. IFN-γ reduced Cy5 signal from 2.84 ± 1.49 to 2.09 ± 1.49 under 100% Trp and from 1.76 ± 1.18 to 0.66 ± 0.65 under 50% Trp. Similarly, Trp deprivation caused a notable decline in Cy5 intensity in both IFN-γ-untreated (2.84 → 1.76) and IFN-γ–treated cells (2.09 → 0.66). These results indicate that both inflammatory and nutrient stress negatively impact the cellular delivery/accumulation of mRNA cargo, possibly through reduced endocytosis, increased degradation, or altered endosomal retention.

To determine whether the drop in eGFP signal could be solely explained by reduced uptake, or whether a defect in translation efficiency also occurred, we performed correlation analysis between Cy5 and eGFP intensities on a per-cell basis (**Fig. 4d–g**). Each dot in these panels represents a single cell, and the slope of the linear regression (constrained through the origin, Y = aX) reflects the average translational yield per unit mRNA. Under control conditions (**Fig. 4d**), a steep slope indicates efficient conversion of mRNA into protein. However, this slope progressively decreased with the addition of IFN-γ (**Fig. 4e**), Trp deprivation (**Fig. 4f**), or both (**Fig. 4g**), demonstrating a significant decline in translational efficiency per mRNA molecule. Importantly, while reduced Cy5 signal explains part of the eGFP loss, the decreasing slope confirms that a true translation defect exists even when accounting for reduced uptake.

Together, these single-cell data disentangle two mechanistic barriers to efficient mRNA-LNP function in dendritic cells: (1) a reduction in cellular mRNA accumulation, likely due to altered uptake or intracellular trafficking, and (2) an independent suppression of translation efficiency. The additive effects of IFN-γ and Trp deprivation on both parameters highlight how immunometabolic cues in the tumor microenvironment or chronic inflammation may hinder mRNA-based interventions, particularly in professional antigen-presenting cells. These results set the stage for further investigation of subcellular trafficking, to be shown in **Fig. 5**.

### Divergent Effects of IFN-γ and Tryptophan Deprivation on Subcellular mRNA Localization, Translation, and Endosomal Escape Efficiency in DC2.4 Dendritic Cells

To interrogate how metabolic and inflammatory cues differentially regulate the intracellular fate of delivered mRNA, we performed spatially resolved, high-resolution confocal imaging and quantitative analysis of mRNA and protein distribution within DC2.4 cells (**Fig. 5a**). Cytoplasmic regions of interest (ROIs) were defined manually for each cell (*n* = 48 per group), excluding the nucleus and lysosomes, to isolate the cytosolic compartment where mRNA translation occurs. For each ROI, we quantified eGFP fluorescence (translated protein) and Cy5 signal (mRNA cargo) per unit area (µm²), enabling direct evaluation of cytoplasmic translation and mRNA localization on a per-cell basis.

Quantification of cytosolic eGFP signal per µm² (**Fig. 5b**) revealed that both tryptophan deprivation and IFN-γ stimulation significantly reduced the translated eGFP in the cytoplasm as the end outcome of mRNA-LNP delivery. Two-way ANOVA identified strong main effects of Trp deprivation (F(1,188) = 92.82, *p* < 0.0001) and IFN-γ (F(1,188) = 33.39, *p* < 0.0001), with no interaction detected, indicating that these stressors independently suppress mRNA-LNP delivery and the final output of eGFP within the cytoplasm. The most profound reduction was observed under the dual treatment condition (50% Trp + IFN-γ), consistent with an additive suppression of mRNA-encoded protein delivery.

To assess whether this reduction in eGFP signal was attributable to lower levels of mRNA available in the cytoplasm, we next quantified cytosolic mRNA-Cy5 signal per µm² (**Fig. 5c**), which reflects the amount of mRNA that successfully escaped the endosomal system and became available in the cytosol. Both Trp deprivation and IFN-γ significantly reduced cytosolic mRNA signal (F(1,188) = 48.52, *p* < 0.0001 and F(1,188) = 26.51, p = 0.0003, respectively), and a significant interaction between them was detected (F(1,188) = 10.91, *p* = 0.0012), suggesting that the combination of inflammatory and metabolic stress alters mRNA availability in a non-additive manner.

To distinguish mRNA abundance effects from *mRNA-to-protein translational efficiency*, we calculated the cytosolic eGFP-to-Cy5 ratio per µm² as a direct measure of mRNA-to-protein translation efficiency within the cytosol (**Fig. 5d**). Two-way ANOVA revealed a significant main effect of Trp deprivation (F(1,188) = 34.91, *p* < 0.0001), but no effect of IFN-γ (F(1,188) = 1.651, *p* = 0.2004) and no interaction (F(1,188) = 0.00342, *p* = 0.9534). This confirms that Trp deprivation specifically impairs mRNA-to-protein translational efficiency, whereas IFN-γ does not. Under 100% Trp, translation efficiency was reduced from 19.33 to 5.89 with Trp deprivation (−69.5%), and from 16.51 to 2.8 with Trp deprivation under IFN-γ (−83.0%), indicating a consistent, dominant effect of nutrient stress on the cell’s capacity to convert mRNA into protein.

We then quantified mRNA signal localized within lysosomes by identifying LysoTracker-positive compartments and measuring Cy5 signal within these ROIs (**Fig. 5e**). This analysis distinguishes uptake and lysosomal sequestration from cytosolic release. Quantification (**Fig. 5f**) showed that IFN-γ significantly reduced lysosomal mRNA content (F(1,188) = 19.13, *p* < 0.0001), while Trp deprivation had no significant effect (F(1,188) = 0.2461, *p* = 0.6204), and no interaction was detected (F(1,188) = 0.8544, *p* = 0.3565). Post hoc analysis revealed that the most substantial reduction occurred between the 50% Trp ± IFN-γ groups (mean difference = 11.22, p = 0.0014), indicating that IFN-γ compromises mRNA uptake or stability within the lysosomal system, independent of Trp availability.

Finally, to evaluate *endosomal escape efficiency*, we calculated the cytosolic-to-lysosomal Cy5 signal ratio per cell as a proxy metric (**Fig. 5g**). Two-way ANOVA detected a significant main effect of Trp deprivation (F(1,188) = 5.974, *p* = 0.0154), but not of IFN-γ (F(1,188) = 0.130, *p* = 0.7197), and no interaction (F(1,188) = 0.078, p = 0.7800). This means Trp deprivation had a significant impact on the endosomal escape efficiency of the mRNA delivered by LNPs. Although individual pairwise comparisons did not reach statistical significance, the trend suggests that Trp deprivation modestly impairs the efficiency of mRNA escape from the endo-lysosomal system into the cytosol. In contrast, IFN-γ has no detectable effect on the endosomal escape efficiency of mRNA-LNPs.

Taken together, these data reveal mechanistically distinct barriers to mRNA-LNP delivery and translation under different stress conditions. IFN-γ primarily impairs mRNA uptake or lysosomal retention, resulting in reduced mRNA availability while leaving translation per mRNA intact. In contrast, Trp deprivation exerts a dual effect, reducing both endosomal escape efficiency and cytoplasmic translational output per mRNA molecule. These findings provide mechanistic insight into how inflammation and nutrient deprivation distinctly constrain mRNA therapeutic efficacy in dendritic cells.

## Discussion

This study aimed to elucidate how two key immunoregulatory cues, namely IFN-γ and tryptophan (Trp) availability, influence the intracellular behavior and translational output of lipid nanoparticle (LNP)-delivered mRNA in dendritic cells. Our central interest lies not in tumor immunotherapy per se, but in understanding how these immunological cues, known to favor tolerogenic programming (e.g., Treg induction)^21^, affect mRNA-LNP-based delivery systems. IFN-γ is a canonical inducer of indoleamine 2,3-dioxygenase 1 (IDO1), which catalyzes the degradation of Trp into kynurenines, creating a locally immunosuppressive and Trp-depleted microenvironment. This pathway plays a crucial role in promoting immune tolerance in various settings, including maternal–fetal tolerance ^22^, transplantation ^23,24^, and chronic inflammation ^25^. We hypothesized that IFN-γ and/or Trp deprivation, either mimicking or bypassing IDO1 activity, would differentially regulate the uptake, intracellular fate, and translational efficiency of mRNA-LNPs, thereby impacting their utility in tolerogenic applications. Our results reveal that IFN-γ and Trp deprivation independently interfere with mRNA-LNP function but act through distinct mechanisms. Specifically, IFN-γ stimulation primarily affects mRNA-LNP uptake and lysosomal accumulation of mRNA cargo. While Trp deprivation also reduces overall mRNA-LNP uptake, it also suppresses translation downstream of uptake and modestly reduces endosomal escape of mRNA-LNPs.

Initial cytotoxicity assays (**Fig. 1c, 3b**) demonstrated that IFN-γ or Trp deprivation alone did not compromise membrane integrity, nor did mRNA-LNP dosing, as indicated by minimal LDH release. However, their combination induced significant cytotoxicity, confirming a synergistic stress phenotype. Importantly, under individual treatments, membrane integrity was maintained, validating subsequent intracellular analyses as cell-autonomous and not confounded by overt toxicity. This was corroborated by luciferase translation assays normalized to LDH (**Fig. 3c**), which showed a dose-dependent decline in mRNA translation under both cues, consistent with imaging-based quantification of per-cell eGFP expression (**Fig. 4b**).

To determine whether this reduction in eGFP resulted from altered mRNA bioavailability (e.g., cellular uptake, endosomal processing, and endo-lysosomal escape of mRNA-LNPs) or mRNA-to-protein translation suppression, we examined mRNA uptake and its intracellular localization. Total Cy5 signal per cell (**Fig. 4c**) was reduced under both IFN-γ and Trp deprivation, without a statistically significant interaction, indicating independent contributions. This points to a shared phenotype of reduced intracellular mRNA accumulation, albeit potentially through distinct upstream mechanisms.

Critically, however, lysosomal mRNA content (Cy5 intensity/µm² in lysosomal ROIs; **Fig. 5f**) was significantly reduced only under IFN-γ, not Trp deprivation. This distinction suggests that IFN-γ alters/reduces uptake and/or potentially promotes lysosomal degradation of the LNP cargo, consistent with literature indicating IFN-γ-driven remodeling of endocytic pathways ^26^. In contrast, Trp deprivation decreased total Cy5 fluorescence per cell (**Fig. 4c**), possibly due to metabolic stress, although ATP level is not reduced under the tested conditions, potentially due to mitochondrial compensation. However, lysosomal Cy5 levels remained unchanged (**Fig. 5f**) under Trp deprivation, either due to (i) reduced endosomal escape rate or (ii) limited lysosomal degradation (likely due to lysosome competition from autophagy induced by Trp deprivation)^27,28^, thus the concentration of the lysosomal mRNA remained similar to standard conditions and significantly higher than INF-γ stimulated conditions.

To further clarify these mechanisms, we interrogated cytosolic mRNA (Cy5) and protein (eGFP) levels in non-lysosomal ROIs (**Fig. 5b–d**). Cytosolic eGFP levels (**Fig. 5b**) and Cy5 levels (**Fig. 5c**) were both reduced by Trp deprivation and IFN-γ, with additive effects. Importantly, cytosolic translation efficiency (eGFP/Cy5 per µm²; Fig. 5d) was significantly impaired under Trp deprivation alone but unaffected by IFN-γ. This indicates that Trp availability directly modulates ribosomal activity, likely via activation of the GCN2-eIF2α-ATF4 axis ^29,30^, a hallmark of the Integrated Stress Response (ISR). The ISR suppresses cap-dependent translation in response to amino acid limitation through phosphorylation of eIF2α, thereby stalling initiation complexes. Our data strongly support this model, as Trp deprivation reduced protein output per unit cytosolic mRNA, independent of mRNA uptake or escape (**Fig. 5d**). In contrast, IFN-γ did not significantly alter this eGFP/Cy5 ratio (**Fig. 5d**), despite reducing both cytosolic mRNA and total Cy5 intensity (**Fig. 5c** and **4c**). This suggests that IFN-γ does not impair ribosomal translation capacity but instead limits mRNA bioavailability upstream, i.e., through reduced uptake (**Fig. 4c**), altered endosomal trafficking (**Fig. 5f**), or enhanced lysosomal degradation (to be investigated in future studies).

The interpretation is further supported by regression analyses of total eGFP versus Cy5 signal per cell (**Fig. 4d–g**), which demonstrated progressively reduced slopes under IFN-γ, Trp deprivation, and their combination. These slopes reflect mRNA-to-protein translation capacity and incorporate all upstream barriers—uptake, escape, and translation. Their decline thus confirms that both cues impair multiple steps of the delivery cascade, with additive, not redundant, effects. The modest yet significant decrease in endosomal escape efficiency (cytosolic-to-lysosomal Cy5 ratio; **Fig. 5g**) under Trp deprivation further suggests that Trp limitation may alter membrane dynamics or endo-lysosomal maturation. While the absolute magnitude of this escape defect was small, it likely contributes cumulatively to the observed suppression of cytoplasmic eGFP levels (**Fig. 5b**). Taken together, our results delineate distinct mechanistic profiles: IFN-γ reduces cellular mRNA levels by affecting uptake and lysosomal retention, whereas Trp deprivation primarily impacts mRNA-LNP uptake and more importantly, it suppresses translational efficiency and endosomal escape efficiency. Importantly, neither cue alone recapitulates the dual suppression observed under combined treatment, which likely reflects compounded blockade at multiple layers, delivery, escape, and translation.

From a translational perspective, this work provides insight into how immune-modulating environments, particularly those enriched in IDO activity and low Trp, affect mRNA therapeutic performance. While such environments are often viewed as barriers to vaccine efficacy, they may be exploited for applications requiring immune suppression or tolerance, such as gene therapy, allergy desensitization, or autoimmune disease. Our data suggest that LNP design strategies should incorporate context-specific optimization, e.g., escape-promoting lipids in Trp-deprived tissues, or uptake-enhancing ligands in IFN-γ–rich sites. More broadly, coupling tolerogenic payloads with environmental sensing elements or stress-insensitive translation motifs may enable more predictable responses in immunosuppressive settings. Future studies should extend these observations in vivo, evaluate their relevance in primary tolerogenic antigen-presenting cells, and explore whether engineering LNP responsiveness to ISR or endosomal dynamics can improve delivery efficacy in challenging microenvironments.

## Conclusion

This study demonstrates that IFN-γ and tryptophan deprivation disrupt mRNA-LNP functionality through distinct but complementary mechanisms. IFN-γ limits mRNA uptake and lysosomal accumulation, whereas tryptophan deprivation suppresses cytosolic translation efficiency and modestly impairs endosomal escape. These findings highlight the importance of immunometabolic context in determining the intracellular fate and translational output of mRNA therapeutics. Understanding how local immune cues influence mRNA-LNP performance provides a foundation for rationally designing delivery systems tailored for tolerogenic applications, such as immune tolerance induction in autoimmune diseases, transplantation, allergy, or AAV gene therapy.

## Acknowledgement

This work was supported by the National Institute of General Medical Sciences of the National Institutes of Health under Award Number R35GM155362.

