## Supplementary material for "Tryptophan and IFN-γ Differentially Modulate Cellular Uptake, Intracellular Trafficking, and Gene Expression of Messenger RNA-Loaded Lipid Nanoparticles in Dendritic Cells": Online Supporting Information: Ma et al_2025_SI_submitted.pdf

^ Equal Contribution.

### Materials

*Cells*

Dendritic cell line DC2.4 (#SCC142) was purchased from Sigma-Aldrich (USA).

*Base Cell Culture Media for DC2.4 Cells*

Tryptophan-free RPMI-1640 with (#R9940) and tryptophan-free RPMI-1640 without phenol red (#23132) were purchased from Teknova. RPMI-1640 with phenol red medium (#25-506) was purchased from Genesee Scientific. Glutamine-free RPMI1640 medium without phenol red (#32404014) was purchased from Fisher Scientific.

*Cell Culture Media Supplements & Reagents*

Penicillin-streptomycin (#15140163), heat-inactivated fetal bovine serum (#A5256801), HEPES buffer solution (Cat# 15630106), GlutaMAX (# 35050061), TrypLE (#12605010), cell culture grade Dulbecco's phosphate-buffered saline (DPBS), IFN-γ recombinant protein (#315-05-20UG), and certified 70% (v/v) reagent alcohol (#LC222105) were purchased from Fisher Scientific (USA). β-mercaptoethanol (#ES-007-E) was purchased from Sigma-Aldrich (USA).

*Cellular Assays*

Cell counting kit-8 (# NC9261855) was purchased from Fisher Scientific (USA). Thiazolyl Blue/MTT (#HY-15924) was purchased from MedChemExpress. The ONE-Glo™ Luciferase Assay System (#E-6120) and CytoTox96® Non-Radioactive Cytotoxicity Assay kit (LDH assay, #G1780) were purchased from Promega (USA). ATP Assay Kit via Firefly Luciferase (Cat# 28854) was purchased from Cell Signaling (USA).

*Lipids & LNP Formulations*

SM-102 (#BP-25499) was purchased from Broadpharm (USA). Cholesterol (#57-88-5), 1,2-distearoyl-sn-glycero-3-phosphocholine (DSPC) (#NC1538812), 1,2-dimyristoyl-rac-glycero-3-methoxypolyethylene glycol-2000 (DMG-$PEG_{2kDa}$) (#NC1665314), were purchased from Avanti Polar Lipids. Invitrogen™ nuclease-free water (# AM9937), Quanti-it™ Ribogreen Reagent and RNA Assay Kit (#R11490), and absolute ethanol (200 Proof) (#64-17-5) were purchased from Fisher Scientific (USA). Microfluidic Mixer MIX-4 Chip (CHP-MIX-4) was purchased from PreciGenome. Molecular biology grade Triton X-100 (#64-846-650ML) was purchased from Sigma-Aldrich (USA). The RNase-free spray (#10-228) was obtained from Genesee Scientific

(USA). HyPure Water for Injection (WFI) Quality Water (#SH31191LS), BI-SCP – Square cuvette/cell, Plastic (#NC9968046) was purchased from Brookhaven Instruments (USA).

#### *mRNAs*

Cy5 EGFP mRNA (#50-199-8310) and negative/scrambled-eGFP mRNA (#50-252-1577) were purchased from APExBIO (USA). CleanCap® Firefly Luciferase mRNA (5-methoxyuridine) (#L-7202) was obtained from TriLink BioTechnologies (USA).

#### *Confocal Microscopic Studies*

Invitrogen™ LysoTracker™ Red DND-99 (#L7528), Hoechst 33342/Tocris Bioscience (#51-17), Leica Microsystems Immersion Oil for Microscopes (#11944399), and Kimwipes™ Delicate Task Wipers (#06-666) were purchased from Fisher Scientific. μ-Slide 8-well high chamber (# NC1930679) was purchased from ibidi America (USA).

#### *Other Consumables*

ClipTip™ Filtered Pipette Tips-200μL (#94420213), ClipTip™ Filtered Pipette Tips-12.5 μL (#94410043), ClipTip™ Filtered Pipette Tips-1250 μL (#94410813), and ClipTip™ Filtered Pipette Tips-20 μL (#94410218) were obtained from Thermo Scientific (USA). Non-sterile and non-treated 96-well black assay plates (#237108) were obtained from Thermo Fisher Scientific (USA). Sterile 96-well clear flat bottom polystyrene microplates, with Lid (#3300) and sterile TC-treated 96-well white plates (#CLS3610-48EA) were obtained from Corning Life Sciences (USA). TC-treated T25 $cm^2$ flask (#25-205) and T75 $cm^2$ (#25-209), 25 mL serological pipettes sterile (#12-104), 10 mL serological pipettes sterile (#12-106), 5 mL serological pipettes sterile (#12-110), and 2 mL aspirating pipette sterile (#12-180) were obtained from Genesee Scientific (USA). Low-binding snap cap microcentrifuge tubes (#07-200-184) were purchased from Costar (USA).

### Methods

#### *General Cell Culture*

DC2.4 cells were cultured in sterile-filtered RPMI-1640 medium (#25-506, Genesee Scientific, USA) supplemented with L-glutamine, 10% (v/v) fetal bovine serum (FBS), 1 mM sodium pyruvate, 1X HEPES buffer solution, 1% Penicillin–Streptomycin, and 0.0054X β-mercaptoethanol. Cells were maintained in a humidified incubator at 37°C with 5% $CO_2$. For further experiments, different types of RPMI media were used, and their compositions are listed in **Table S1 below**.

#### *LDH Assay*

Cell membrane integrity, indicative of cytotoxicity and indirectly of cell viability, was assessed by measuring lactate dehydrogenase (LDH) release into the culture medium. The supernatant from the untreated DC2.4 cells and the lysed untreated DC2.4 cells were used as reference points for 0% and 100% LDH release, respectively. The measurement of LDH in the supernatant was measured by CytoTox 96® Reagent (Promega) following the manufacturer's instructions. The readout is the absorbance measured at 490 nm using a VarioSkan™ microplate reader.

#### *Mitochondrial activity measured by MTT assay*

DC2.4 cells were seeded at a density of 4,000 cells per well in clear, flat-bottom 96-well plates and incubated for 12 hours at 37 °C in a humidified incubator with 5% $CO_2$. After 12 hours of seeding, the culture medium was replaced with 100% Tryptophan medium/50% tryptophan-deprivation medium with & without IFN-γ–supplementation (100 ng/mL) conditioned. Following 12 hours of exposure to the conditioned media, LNPs encapsulating mRNA with a dose of 100 ng Fluc mRNA per well were added to the cells in the continued presence of the same media. MTT reagent was prepared by dissolving Thiazolyl Blue Tetrazolium Bromide in 1X Dulbecco's phosphate-buffered saline (DPBS) at a concentration of 5.0 mg/mL (10X stock), protected from light, and stored at 4 °C for further use. After a 24-hour incubation with LNPs, the treatment media were carefully aspirated. To each well, 90 µL of pre-warmed complete RPMI-1640 medium was added along with 10 µL of the 5.0 mg/mL MTT stock solution, yielding a final MTT concentration of 0.5 mg/mL. Plates were then incubated for 4 hours at 37 °C, protected from light. After incubation, the MTT-containing medium was gently removed, and 100 µL of dimethyl sulfoxide (DMSO) was added to each well to dissolve the formazan crystals. Plates were placed on an orbital shaker at 300 rpm for 15 minutes to ensure complete solubilization. Absorbance was measured at 570 nm using a VarioSkan™ microplate reader. GraphPad Prism 10 was used for graphical presentation and statistic calculation.

#### *Overall cellular metabolic activity measured CCK-8 assay*

Overall cellular metabolic activity was evaluated using the Cell Counting Kit-8 (CCK-8; Dojindo Molecular Technologies) according to the manufacturer's instructions. DC2.4 cells were seeded at a density of 4,000 cells per well in clear, flat-bottom 96-well plates and incubated for 12 hours at 37 °C in a humidified incubator with 5% $CO_2$. After 12 hours, the culture medium was replaced with 100% Tryptophan medium/50% tryptophan-deprivation medium with & without IFN-γ-supplementation (100 ng/mL) conditioned. Following a 12-hour pre-treatment period, LNPs encapsulating mRNA with a dose of 100 ng Fluc mRNA per well were added to the cells in the same conditioned media. After a subsequent 24-hour incubation with LNPs, all treatment media were replenished with 90 µL of pre-warmed RPMI1640 complete culture medium, and 10 µL of CCK-8 reagent was added directly to each well containing 100 µL of culture medium, yielding a final 1:10 dilution. Plates were incubated for 4 hours at 37 °C, protected from light. Absorbance was measured at 450 nm using a VarioSkan™ microplate reader. Cell viability was calculated relative to untreated control wells after background correction. GraphPad Prism 10 was used for graphical presentation and statistical calculation.

#### *Cellular ATP production validated by firefly luciferase ATP assay*

Cellular ATP production was assessed using the Firefly Luciferase ATP Assay Kit (Cell signaling, USA) to validate metabolic activity following treatment. DC2.4 cells were seeded at a density of 4,000 cells per well in white, opaque 96-well plates and incubated for 12 hours at 37 °C with 5% $CO_2$. After 12h of post-seeding, the culture medium was replaced with one of the following treatment conditions: 100% Tryptophan medium/50% tryptophan-deprivation medium with & without IFN-γ–supplementation (100 ng/mL) conditioned. After a 12-hour pre-treatment period, mRNA-loaded LNPs with a dose of 100 ng Fluc mRNA per well, were added to the cells in the respective media. Following a 24-hour incubation with LNPs, all conditioned media were replenished with 100 µL of complete RPMI1640 cell culture media and 100 µL of the FLuc substrate reagent was added to each well according to the manufacturer's instructions, and luminescence was measured using a VarioSkan™ microplate reader. ATP levels were quantified relative to untreated controls after background subtraction. GraphPad Prism 10 was used for graphical presentation and statistic calculation.

#### *mRNA-Lipid Nanoparticle (LNP) Preparation and Characterization*

The LNP was prepared using the rapid ethanol injection method reported previously [1,2]. In brief, lipid formulation was prepared in ethanol with a mixture of SM-102, 1,2-distearoyl-sn-glycero-3-phosphocholine (DSPC), cholesterol, and 1,2-dimyristoyl-rac-glycero-3-methoxypolyethylene glycol-2000 (DMG-$PEG_{2kDa}$) at a molar ratio of 50:10:38.5:1.5. The formulation solution was then mixed with mRNA solution in citric acid buffer (pH 4.0) at an NP ratio of 1:6 and a volume ratio of 1: 3 (v/v) following in-line dilution and stored at 4 °C before use. The mRNA encapsulation efficiency (% mRNA recovery) was evaluated using the Quant-iT RiboGreen RNA Assay Kit, following the manufacturer's protocol. Molecular biology grade Triton X-100 was used to lyse the LNPs for mRNA quantification. Fluorescence measurements were obtained with a VarioSkan™ LUX microplate reader (Thermo Fisher Scientific) using an excitation wavelength of 480 nm and an emission wavelength of 520 nm. The EE% is calculated using (Fluorescence of formulated mRNA-LNPs)/(Fluorescence of Triton X-100 lysed mRNA-LNPs) x 100%. The hydrodynamic diameter and zeta potential of LNPs were characterized using a NanoBrook 90Plus PALS instrument (Brookhaven Instruments, USA).

#### *Protein Expression Screening using mRNA-FLuc Luciferase Reporter*

DC2.4 cells were seeded at a density of 4,000 cells per well in white 96-well plates and incubated at 37°C with 5% $CO_2$. After 12h incubation, the culture medium was replaced with four different RPMI-1640 media: 1) standard media with 100% tryptophan medium, with and without IFN-γ (100 ng/mL); 2) 50% tryptophan-deprivation medium, with and without IFN-γ (100 ng/mL), and then incubated for another 12h. The media were then aspirated and replaced with 200 µL/well of conditioned medium containing mRNA-LNPs encoding firefly luciferase (FLuc), at a final mRNA dose of 200 ng per well. Cells were incubated for an additional 24 hours to allow for FLuc translation from the mRNA. After the incubation, the supernatant was removed for the downstream LDH assay. The FLuc bioluminescence was quantified by adding ONE-Glo™ Luciferase reagent (100 µL/well, incubated for 2 minutes), and then measured using a VarioSkan™ LUX microplate reader (Thermo Fisher Scientific). The luminescence signal was then normalized by cell viability % from the LDH assay.

#### *Confocal microscopy studies*

To quantify antigen mRNA delivery and translation efficiency via LNPs under cellular stressed or essential amino acid deprivation condition, live cell confocal microscopy imaging was performed. DC2.4 cells (30,000 cells per well) were seeded into µ-Slide 8-Well high chambers (ibidi GmbH, Germany) and incubated for 12 hours at 37°C with 5% $CO_2$. The culture medium was then replaced with one of the following treatment conditions: 100% Tryptophan medium, 50% tryptophan-deprivation medium, with or without IFN-γ supplementation (100 ng/mL), conditioned. After a 12-hour pre-treatment period, mRNA-loaded LNPs encapsulating 200 ng of eGFP-Cy5 mRNA were added to each well in the respective treatment media. Following 24 hours of incubation, the culture medium was removed, and live-cell staining was performed. Cells were incubated with 50 nM LysoTracker™ Red DND-99 (Thermo Fisher Scientific) for 1 hour, followed by staining with 5 µg/mL Hoechst 33342 (Thermo Fisher Scientific) for 10 minutes. After staining, fresh medium was added, and images were acquired using a Leica TCS SP8 confocal microscope (Leica Microsystems, Germany), equipped with a 63X/1.4 NA oil-immersion objective. Images were processed and exported using LAS X software. Fluorescence quantification was measured by ImageJ v1.5.4p software (NIH, USA). GraphPad Prism 10 was used for graphical presentation and statistical calculation.

**Table S1: Different RPMI 1640 media compositions with & without Phenol Red and with & without Tryptophan**

| Media Composition | With Tryptophan & with Phenol Red Media (RPMI1640, #25-506, Genesee Scientific) | Without Tryptophan & with Phenol Red Media (RPMI1640, #R9940 Teknova) | With Tryptophan & without Phenol Red Media (RPMI1640, #32404-014, Gibco) | Without Tryptophan & without Phenol Red Media (RPMI1640, #23132, Teknova) | With Tryptophan & with Phenol Red Media (RPMI1640, #10-040-CV, Corning) |
|---|---|---|---|---|---|
| RPMI1640 | 480mL | 480 mL | 480 mL | 480 mL | 475 mL |
| Fetal bovine serum | 5 mL | 5 mL | 5 mL | 5 mL | 5 mL |
| HEPES Buffer | 5 mL | 5 mL | 5 mL | 5 mL | 5 mL |
| Glutamax | - | 5 mL | 5 mL | 5mL | - |
| Penicillin/Streptomycin | 5 mL | 5 mL | 5 mL | 5 mL | 5 mL |
| DPBS | 5 mL | - | - | - | 5 mL |
| Sodium Pyruvate | - | - | - | - | 5 mL |
| Total Volume | 500 mL | 500 mL | 500 mL | 500 mL | 500 mL |
